# Synaptic Basis of Behavioral Timescale Plasticity

**DOI:** 10.1101/2023.10.04.560848

**Authors:** Kevin C. Gonzalez, Adrian Negrean, Zhenrui Liao, Franck Polleux, Attila Losonczy

## Abstract

Learning and memory are fundamental to adaptive behavior and cognition. Various forms of synaptic plasticity have been proposed as cellular substrates for the emergence of feature selectivity in neurons underlying episodic memory. However, despite decades of work, our understanding of how synaptic plasticity underlies memory encoding remains limited, largely due to a shortage of tools and technical challenges associated with the visualization of synaptic plasticity at single-neuron resolution in awake-behaving animals. Behavioral Timescale Synaptic Plasticity (BTSP) postulates that synaptic inputs active during a seconds-long time window preceding and immediately following a large depolarizing plateau spike are potentiated, while synaptic inputs active outside this time window are depressed. We experimentally tested this model *in vivo* in awake-behaving mice using an all-optical approach by inducing place fields (PFs) in single CA1 pyramidal neurons (CA1PNs) while monitoring the spatiotemporal tuning of individual dendritic spines and changes in their corresponding synaptic weights. We identified an asymmetric synaptic plasticity kernel resulting from bidirectional modifications of synaptic weights around plateau burst induction. Surprisingly, our work also uncovered compartment-specific differences in the magnitude and temporal expression of synaptic plasticity between basal and oblique dendrites of CA1PNs. Our results provide the first experimental evidence linking synaptic plasticity to the rapid emergence of spatial selectivity in hippocampal neurons, a critical prerequisite for episodic memory.

---

The mammalian brain learns continuously throughout an individual’s lifetime, with an astonishing capacity to acquire, retain and retrieve relevant new information while simultaneously filtering and forgetting irrelevant experiences. The primary neural basis for these behavioral adaptations is thought to be synaptic plasticity, which underlies changes in the functional connectivity of neuronal circuits in the brain^1^. Various forms of experience-dependent synaptic modifications^2^, particularly at excitatory glutamatergic synapses, are widely considered to be the primary substrate of learning^3–5^. However, causal links have yet to be made between the rules of synaptic plasticity and memory formation due to the difficulty of monitoring and manipulating plasticity at the synaptic scale *in vivo*. In the hippocampus, individual excitatory CA1PNs form PFs during exploratory behavior^6^, a phenomenon in which a neuron will fire action potentials only in a restricted sub-region of physical space. In these neurons, experience-dependent spatial tuning is thought to reflect the formation of a malleable conjunctive representation of space and contextual valence, linking a spatial representation of the environment with the sensory details of the associated experience^7–9^. While the functional properties of hippocampal circuits supporting spatial and episodic memory^10–12^ are traditionally investigated and interpreted at the level of place cells^6,13–16^, the synaptic mechanisms underlying PF formation have remained largely unknown. Recently, BTSP has been identified as a prevalent form of plasticity underlying PF formation in hippocampal CA1PNs^16,17^. This form of plasticity leads to the rapid, single-trial formation of PFs following spontaneous dendritic plateau-burst spikes, as well as following electrophysiological or optogenetic induction^16–22^. *In vitro* recordings^16,17^ and recent modeling^23^ suggest that a seconds-long, asymmetric synaptic plasticity kernel underlies BTSP due to both potentiation of glutamatergic synapses active several seconds before and during the plateau-burst events, and depression of synaptic inputs active outside this temporal window. However, the synaptic rules associated with BTSP have not been directly measured *in vivo* in awake-behaving animals during spatial navigation. Here, we developed a single-cell-based, all-optical approach to simultaneously monitor the spatial tuning of individual inputs as well as changes in their synaptic weights before and after PF induction to uncover the synaptic plasticity rules underlying BTSP in hippocampal CA1PNs.

## *In vivo* functional imaging of dendritic spine activity in hippocampal region CA1

To investigate the relationship between changes in synaptic weight and the emergence of PFs in individual CA1PNs, we developed and deployed three tools at single-cell resolution (Fig. 1a): (1) To measure the spatial tuning of excitatory synaptic inputs received by dendritic spines and the timing of their activity relative to an experimentally induced dendritic plateau potential, we used a genetically encoded glutamate release sensor (SFVenus-iGluSnFr-A184S). This sensor, when expressed postsynaptically, reports presynaptic glutamate release of individual inputs received by dendritic spines^24–26^; (2) To measure changes in functional synaptic strength (synaptic weight, *W*) *in vivo* before and after PF induction, we used a red-shifted calcium indicator (jRGECO1a) to monitor spine calcium dynamics. We focused on spine calcium because their levels rise in response to synaptic activity. This increase is mediated by *N*-methyl-D-aspartate receptors (NMDARs) and voltage-gated calcium channels (VGCCs)^27,28^, and is dependent on the degree of synaptic depolarization provided by AMPA receptors (AMPARs)^29–33^; (3) To induce PFs, we used a red-shifted excitatory opsin (bReaChes). The spatially restricted optogenetic activation of bReaChes expressing cells can effectively generate long-lasting PFs^18–21^.

**Fig 1:**
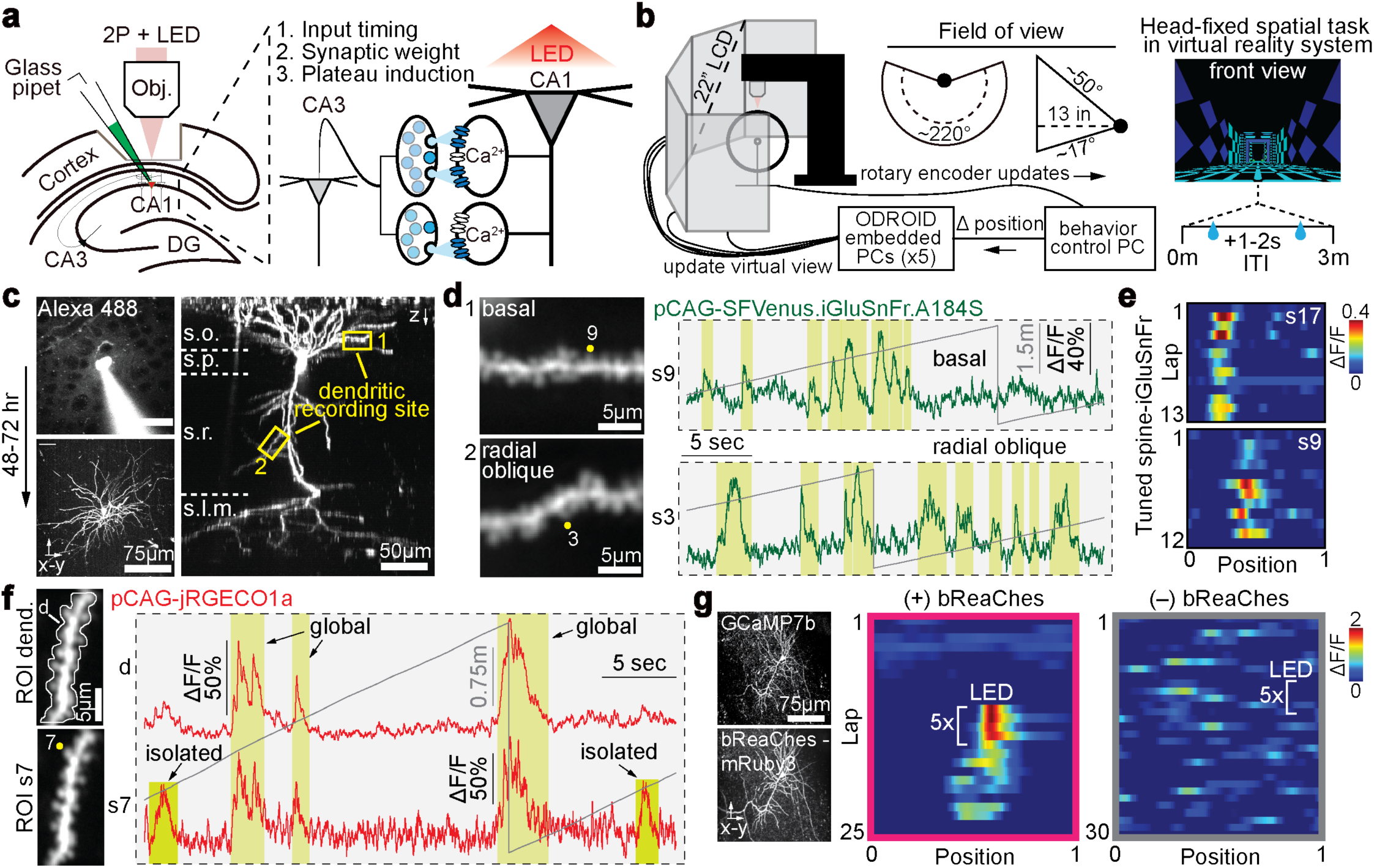
*In vivo* subcellular resolution imaging of presynaptic glutamate release and postsynaptic spine calcium activity before and after optogenetic place field induction in single CA1 pyramidal neurons. **a**, Schematic of *in vivo* two-photon (2P) guided single-cell electroporation (SCE) setup and proposed functional readout of electroporated plasmids. Cells express a depolarizing opsin (bReaChes) to induce place fields and two sensors to measure input timing (using the postsynaptic membrane-bound glutamate sensor iGluSnFr) and changes in synaptic weight (using the postsynaptic cytosolic calcium sensor jRGECO1a). **b**, Schematic of combined 2P and virtual reality system. Mice perform a head-restricted spatial navigation task for fixed water rewards. ITI, intertrial interval. **c**, Glass pipets containing plasmids and Alexa488 were used to guide the electroporation of single cells in the pyramidal layer (top left). Cells reached peak expression 36–48 hours post-SCE (bottom left). Dendrites spanning the entire dendritic tree were accessible (right). s.o., stratum oriens; s.p., stratum pyramidale; s.r., stratum radiatum; s.l.m., stratum lacunosum moleculare. **d**, Increased magnification of the areas indicated in **c** showing maximum intensity projections of single-plane time-series imaging of iGluSnFR-expressing basal and radial oblique dendrites, along with associated fluorescence traces of a subset of spines (yellow circles). Gray traces represent the animal’s position on the virtual track. **e**, Synaptic activity heatmaps for two spines receiving spatially-tuned iGluSnFr input. **f**, Example maximum intensity projection of a single-plane time-series imaging of a jRGECO1a-expressing basal dendrite. Associated fluorescence traces acquired from the dendritic (white outline) and synaptic (yellow circle) regions of interest are shown. Gray traces represent the animal’s position on the virtual track. **g**, Single pyramidal cell expressing a red-shifted excitatory opsin (bReaChes-mRuby3) and GCaMP7b. Spatially restricted optogenetic stimulations (LED) for five consecutive laps evoke strong somatic responses and induce long-lasting place fields (+)(magenta). These effects are lost in cells electroporated without the opsin plasmid (–)(gray).

With these tools, we performed *in vivo* single-cell electroporation (SCE)^19–21^ to introduce plasmids into individual pyramidal neurons located within the dorsal CA1 region of the mouse hippocampus. As described above, we introduced the genetically encoded glutamate release sensor (pCAG-SFVenus-iGluSnFr-A184S) and the calcium indicator (pCAG-jRGECO1a) to estimate the spatial tuning properties of inputs received by individual dendritic spines and changes in synaptic weight following an experimentally evoked plateau potential produced by the excitatory opsin, bReaChes (pCAG-bReaChes-mRuby3)^21^. Using a head-fixed spatial navigation task in a 3 meter-long virtual reality (VR) environment^34,35^, we carried out two-photon (2P) imaging of synaptic activity dynamics from basal (*stratum oriens*, SO) and oblique (*stratum radiatum*, SR) dendrites, which receive the majority of spatially tuned inputs from upstream intrahippocampal regions CA2/CA3 and are thought to be the primary inputs acted upon by rapid plasticity mechanisms to generate PFs^23^ (Fig. 1b, c). Using this approach, we confirmed that we could reliably detect stable, spatially tuned synaptic inputs using the glutamate sensor iGluSnFr (Fig. 1d, e, Extended Data Fig. 1a–e) and that we could distinguish action potential-related global dendritic events from isolated spine-specific activity at individual spines using the calcium indicator jRGECO1a (Fig. 1f). Lastly, in line with previous results, we showed that we could perform optogenetic stimulation of bReaChes to induce PFs that exhibit characteristic features of endogenously occurring PFs formed through BTSP *in vivo* (Fig. 1g, Fig. 2b).

**Fig 2:**
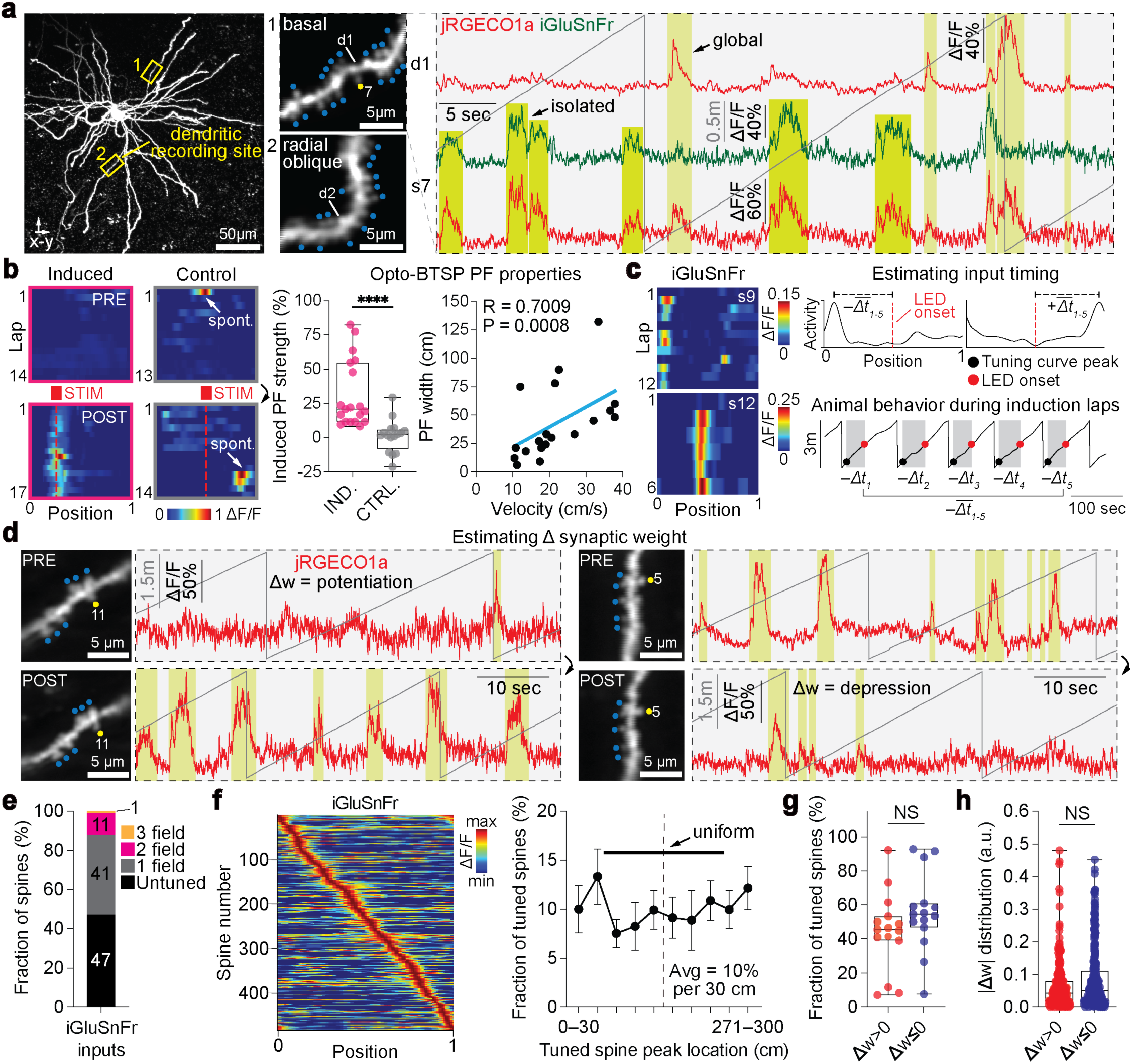
Optical measurement of synaptic plasticity associated with induction of hippocampal BTSP. **a**, *In vivo* images (maximum intensity projections) of jRGECO1a, iGluSnFr, and bReaChes expressing cells. Increased magnification of recorded dendritic segments and associated fluorescence traces of synaptic glutamate–calcium (yellow circle) and dendritic calcium signals during navigation are shown. Laterally protruding spines are labeled with blue circles. Gray traces represent the animal’s position on the virtual track. **b**, Left: Before (PRE) and after (POST) somatic activity heatmaps for induced (magenta) and control (gray) groups. Red boxes indicate the stimulation site. Red dashed lines indicate the expected induced place field (PF) location. White arrows show spontaneous PF formation events in the control group. Center: Optogenetic stimulation significantly increases somatic activity within the LED zone (see methods) (Mann-Whitney two-tailed unpaired *t*-test: *n* = 19 induction experiments (12 cells), *n* = 16 control experiments (11 cells), *P* < 0.0001). Right: Strong relationship between experimentally induced PF width and velocity on the first induction lap (Spearman r: *n* = 19 induction experiments (12 cells), two-tailed *t*-test, *R* = 0.7009, *P* = 0.0008). Data points fit by linear equation (blue line). **c**, Activity heatmaps for two spines receiving spatially tuned iGluSnFr input and schematic illustrating the extraction of input timing. The average time between the peak of the detected spine PF and the LED onset is measured using the animal’s behavior during the five induction laps. Positive time values mean the input arrived after the LED onset. Negative time values mean the input arrived before the LED onset. **d**, *In vivo* images of dendrites recorded before (PRE) and after (POST) place field induction, along with associated synaptic calcium signals for a subset of spines undergoing potentiation and depression (yellow circles). To measure changes in synaptic weight, the amplitude of all isolated spine-specific calcium events in the trace were summed and compared (POST–PRE) (see methods). Gray traces represent the animal’s position on the virtual track. **e**, Fraction of spines (*n* = 906 spines total) receiving spatially tuned synaptic iGluSnFr input to one (gray), two (magenta), or three (orange) locations, and spines receiving no tuned input (black). **f**, Left: average iGluSnFr ΔF/F across space for all tuned spines sorted by peak location (*n* = 484 spines from 15 induction experiments (12 cells)). Right: fraction of tuned spines versus PF peak location (bin = 30 cm). Data are shown as mean ± s.e.m. Excitatory inputs are homogeneously distributed in the environment (chi-square test of independence, df = 9, *P* = 0.451). **g**, Fraction of tuned spines undergoing potentiation or depression per experiment (Mann-Whitney two-tailed unpaired *t*-test: *n* = 15 induction experiments (12 cells); *P* = 0.0998). **h**, Magnitude of potentiation and depression events occurring at tuned spines (Mann-Whitney two-tailed unpaired *t*-test: *Δw*>0, *n* = 213 spines; *Δw*≤0, *n* = 271 spines; *P* = 0.0810). All box plots depict the median (central line) and interquartile range (25^th^ and 75^th^ percentile). Whiskers extend to the min-max data points.

## Measuring spatial tuning of excitatory synaptic inputs and changes in synaptic weights before and after optogenetic place field induction

We next combined our single-cell labeling approach with optogenetic manipulations to measure the synaptic plasticity (i.e., changes in synaptic weight, *ΔW*) associated with the induction of a PF. To achieve this, we co-electroporated the glutamate and calcium sensors and a modified, untagged version of the bReaChes opsin lacking the mRuby3 fluorophore. This allowed us to simultaneously perform dendritic calcium imaging and functional input mapping before and after PF induction (Fig. 2a, b).

To measure the timing of synaptic input arrival relative to the predetermined location of PF induction while also monitoring changes in synaptic weight, we divided our experiments into three phases: *Pre*, *Induction*, and *Post*. In the *Pre* phase, we sampled the tuning specificity of the soma, mapped the spatiotemporal distribution of synaptic inputs, and measured the initial synaptic weights of dendritic spines before induction (Fig. 2b-d). In the *Induction* phase, we induced a PF (Fig. 2b). In the *Post* phase, we confirmed the formation of a PF and measured the final weights of all spines after induction (Fig. 2b, d). We restricted our analysis to spines receiving tuned inputs in order to estimate the time interval between the arrival of the input and the site of optogenetic stimulation (Fig. 2c). To measure plasticity, we focused exclusively on the properties of spine-restricted calcium signals, which arise from synaptic activation of NMDARs and VGCCs^27,28,32,33^. Since plasticity can influence either the amplitude or frequency of the detected spine-specific calcium events, we quantified synaptic weight by summing the amplitudes of these events, using this measurement as an indicator of both aspects of plasticity. To ensure that the plasticity measurements were not contaminated by the expected changes in somatic firing properties following PF formation, we excluded all spine calcium events that co-occurred with globally detected dendritic calcium events (Fig. 1f, 2a, see methods).

In total, we performed 19 optogenetic inductions from a set of CA1PNs (12 cells). In about half of these recordings, the neurons had a preexisting somatic PF during the *PRE* phase of the experiment (9/19). We imaged synaptic activity from 71 dendritic branches (40 basal, 31 oblique, 906 spines total). Approximately half of the dendritic spines we imaged received stable, spatially tuned, excitatory synaptic input (Fig. 2e) that evenly tiled the VR environment (Fig. 2f). On a per cell basis, we observed a broad distribution in the number of tuned spines undergoing potentiation and depression (Fig. 2g), and on a per spine basis, a broad distribution in the magnitude and direction of the plasticity (Fig. 2h). We observed no significant difference in the distribution or amplitude of the plasticity events between tuned or untuned spines in both the induced (with opsin) and control (without opsin) groups (Extended Data Fig. 2a–b), suggesting that background synaptic plasticity occurs even without optogenetic stimulation. In line with this, we observed multiple spontaneous remapping events in our *Pre* and *Post* somatic control traces closely resembling BTSP-like PF formation events (Fig. 2b). However, we did find that large magnitude synaptic plasticity events (upper 25^th^ percentile of all *|ΔW|*) were selectively evoked at tuned compared to untuned spines in the induced group (Extended Data Fig. 3a), suggesting that spines are filtered to undergo synaptic plasticity based on the tuning profile of the inputs they receive.

## Spatial and temporal profiles of optically induced BTSP

Next, we characterized how the measured synaptic plasticity events were temporally coordinated around the time of PF induction. To do so, we focused on the asymmetric 6-second time window (4 seconds pre-stimulation to 2 seconds post-stimulation), where BTSP has been hypothesized to exert its most pronounced effects^16,23^. We found that plasticity was strongly and inversely correlated with the initial synaptic weight of each synapse (Fig. 3a) and highly coordinated in time, resulting in a kernel of synaptic depression before (∼3-4 seconds) and after (∼1-2 seconds) the induction, and synaptic potentiation immediately before (∼1-2 seconds) the induction (Fig. 3b, left). The peak of the temporal kernel not only occurred before the stimulus but also decayed asymmetrically in time (Fig. 3b, left: ρ_b_ (tau backward) from single exponential fit of data ranging from - 4.0 to -1.0 seconds (ρ_b_ = 2.00 seconds; R^2^ = 0.9447). ρ_f_ (tau forward) from single exponential fit of data ranging from -1.0 to +2.0 seconds (ρ_f_ = 0.60 seconds; R^2^ = 0.8736). This temporal kernel became even more pronounced when we restricted our analysis to spines undergoing significant changes in synaptic weights (upper 25^th^ percentile of all *|ΔW|*) (Fig. 3b, center) and was completely absent in control CA1PNs that received LED stimulation but were not expressing any opsin (Fig. 3b, right). Moreover, this plasticity was not accompanied by any detectable differences in spine head size between potentiated and depressed spines (Extended Data Fig. 6a–b).

**Fig 3:**
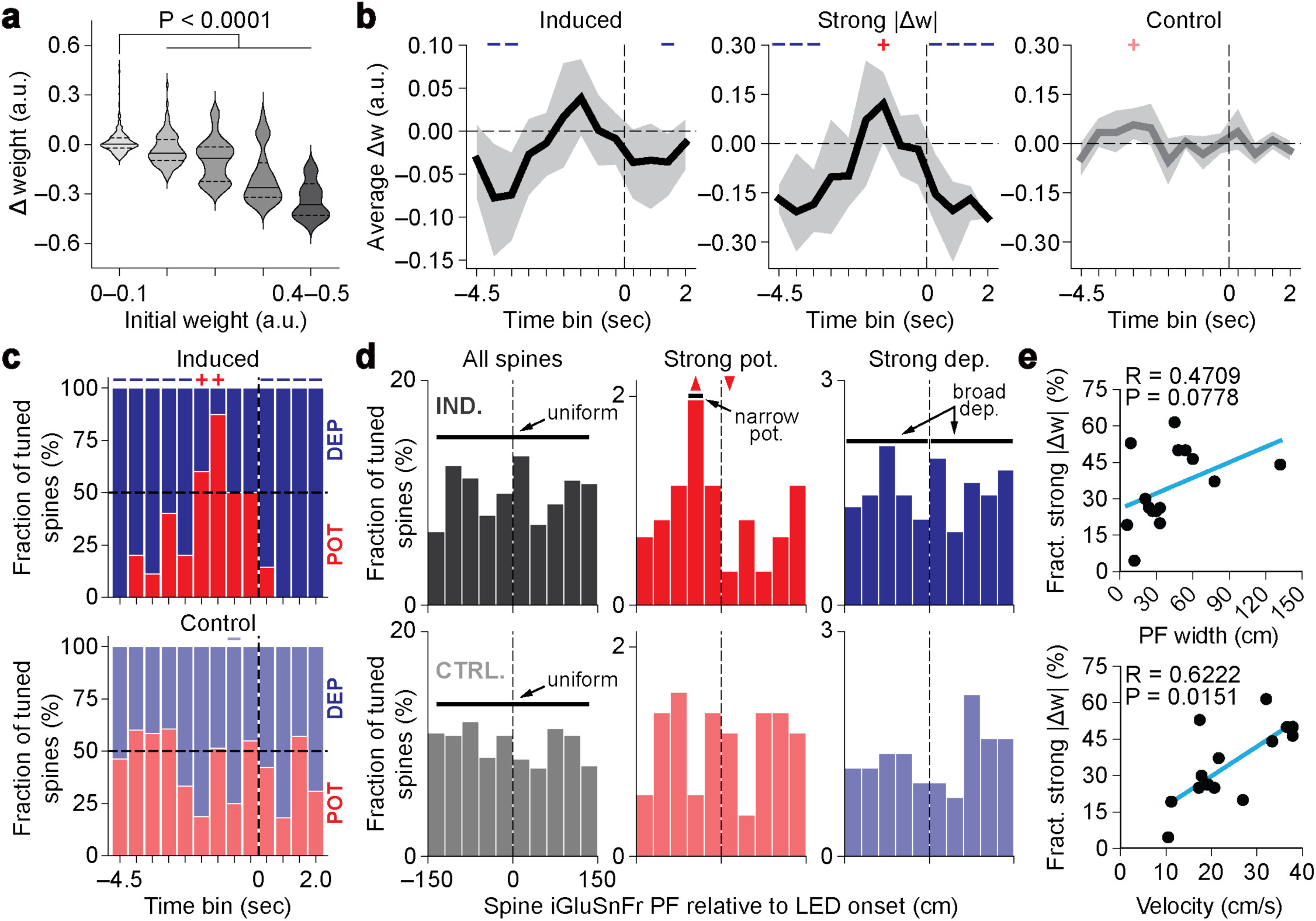
Coordinated bidirectional changes in synaptic weight underlie behavioral timescale plasticity. **a**, Magnitude of synaptic plasticity events (Δ weight) versus initial weight (bin = 0.1 a.u., Kruskal-Wallis tests, solid lines show the median, dashed lines show the 25^th^ and 75^th^ percentiles). **b**, Temporal profile of average plasticity occurring around the time of optogenetic place field (PF) induction (bin = 0.5 seconds; left: induced group, *n* = 226 spines; center: same as induced, except only looking at the upper 25^th^ percentile of all plasticity events occurring at tuned spines, *n* = 62 spines; right: control group, *n* = 224 spines). Blue minus symbols depict time bins in which depression dominates and red plus symbols depict those in which potentiation dominates (bootstrapped 95% confidence intervals (CIs) were generated from 10,000 bootstraps of the data points in each time bin; significance was assigned to time bins where CIs did not cover zero). **c**, Temporal profile of fraction of spines undergoing potentiation and depression around the time of optogenetic PF induction (bin = 0.5 seconds; top: induced group, *n* = 62 spines (see Fig. 3b center); bottom: control group, *n* = 224 spines (see Fig. 3b right). The blue minus (significant depression) and red plus (significant potentiation) symbols indicate time bins that lie outside the 95% CI generated from 1,000 data shuffles. **d**, Spatial profile of synaptic plasticity events (top row = induced group; bottom row = control group; bin = 30cm). Left: excitatory inputs are homogenously distributed around the location of optogenetic PF induction (chi-square test of independence, df = 9; IND., *P* = 0.114; CTRL., *P* = 0.804). Center: spines receiving excitatory inputs tuned to locations before the stimulation site are selectively recruited to undergo strong potentiation, while spines receiving excitatory inputs tuned to locations after the stimulation site are actively suppressed from undergoing potentiation. Right: spines are broadly depressed. Data were thresholded only to include the upper 25^th^ percentile of all plasticity events occurring at tuned spines. The up and down arrowheads indicate position bins that lie above or below the 95% CI generated from 1,000 data shuffles, respectively. **e**, Fraction of tuned spines recruited to undergo strong plasticity around the time of optogenetic PF induction as a function of PF width (top; Spearman r: *n* = 15 induction experiments (12 cells), two-tailed *t*-test, *R* = 0.4709, *P* = 0.0778) and animal velocity (bottom; Spearman r: *n* = 15 induction experiments (12 cells), two-tailed *t*-test, *R* = 0.6222, *P* = 0.0151). Fraction was calculated by dividing the number of tuned spines undergoing strong plasticity inside the kernel (−4.5 to +2.0 seconds) by the total number of tuned spines recorded in the experiment. Spines undergoing no plasticity (bottom 25^th^ percentile of all plasticity events occurring at tuned spines) were removed from the analysis. Data are fitted by a linear equation (blue line).

We corroborated these results by looking at the fraction of spines undergoing potentiation and depression as a function of time. Using this metric, which is independent of the amplitude of each plasticity event, we found that the inputs active immediately prior to the stimulation site preferentially underwent potentiation, while the inputs active >2 seconds before or >0.5 seconds after optogenetic induction tended to experience depression (Fig. 3c, top). This temporal kernel was absent in control cells that received LED stimulation but were not expressing any opsin (Fig. 3c, bottom).

We next determined the spatial profile of the induced plasticity. We observed a uniform distribution of excitatory inputs active around the induction site during the *Pre* phase of the experiment (Fig. 3d, left). However, following PF formation, we found that spines receiving inputs active when the animal was located before the induction site underwent significant potentiation, while spines receiving inputs active when the animal was located after the induction site were actively suppressed from undergoing potentiation. (Fig. 3d, center). This site-specific potentiation was accompanied by a broad depression of inputs tuned to all spatial locations (Fig. 3d, right). Lastly, we examined the relationship between PF width and velocity on the fraction of spines recruited to undergo plasticity. Corroborating previous work^16^, we found that both parameters correlated strongly with the fraction and spatial coverage of the inputs undergoing plasticity (Fig. 3e). Together, these results demonstrate that the emergence of a PF in CA1PNs is associated with a temporally structured, bidirectional kernel of changes in synaptic weights organized along a seconds-long timescale.

## Compartmentalized expression of synaptic plasticity following place cell induction

Recent studies have revealed that basal and apical oblique dendrites of CA1PNs have distinct synaptic organization and functional properties^21,36–39^. We took advantage of the fact that we recorded dendritic spine activity from basal and oblique dendrites to test whether the spatiotemporal rules of synaptic plasticity we described above apply equally to both dendritic compartments. We found that oblique spines received significantly less spatially tuned excitatory inputs (Extended Data Fig. 5a) and experienced moderately different degrees of synaptic potentiation and depression compared to basal spines (Fig. 4a). To investigate this further, we compared the initial synaptic weight distributions in both compartments and found that oblique spines displayed significantly higher initial weights than basal spines (Fig. 4b). Based on our previous results showing a negative correlation between initial synaptic weight and propensity to undergo larger changes in synaptic weight (Fig. 3a), this result suggested that spines located along oblique dendrites could be more readily recruited to undergo bidirectional changes in plasticity compared to basal spines. Indeed, when we measured the amplitude and direction (potentiation vs. depression) of the changes in synaptic weights, we found that oblique spines exhibited significantly larger changes in synaptic weights and underwent more potentiation and depression compared to basal spines (Fig. 4c, Extended Data Fig. 5c, d). Moreover, when we compared the cumulative plasticity in each dendritic compartment, we observed that oblique spines underwent significantly more depression than basal spines (inset in Fig. 4c).

**Fig 4:**
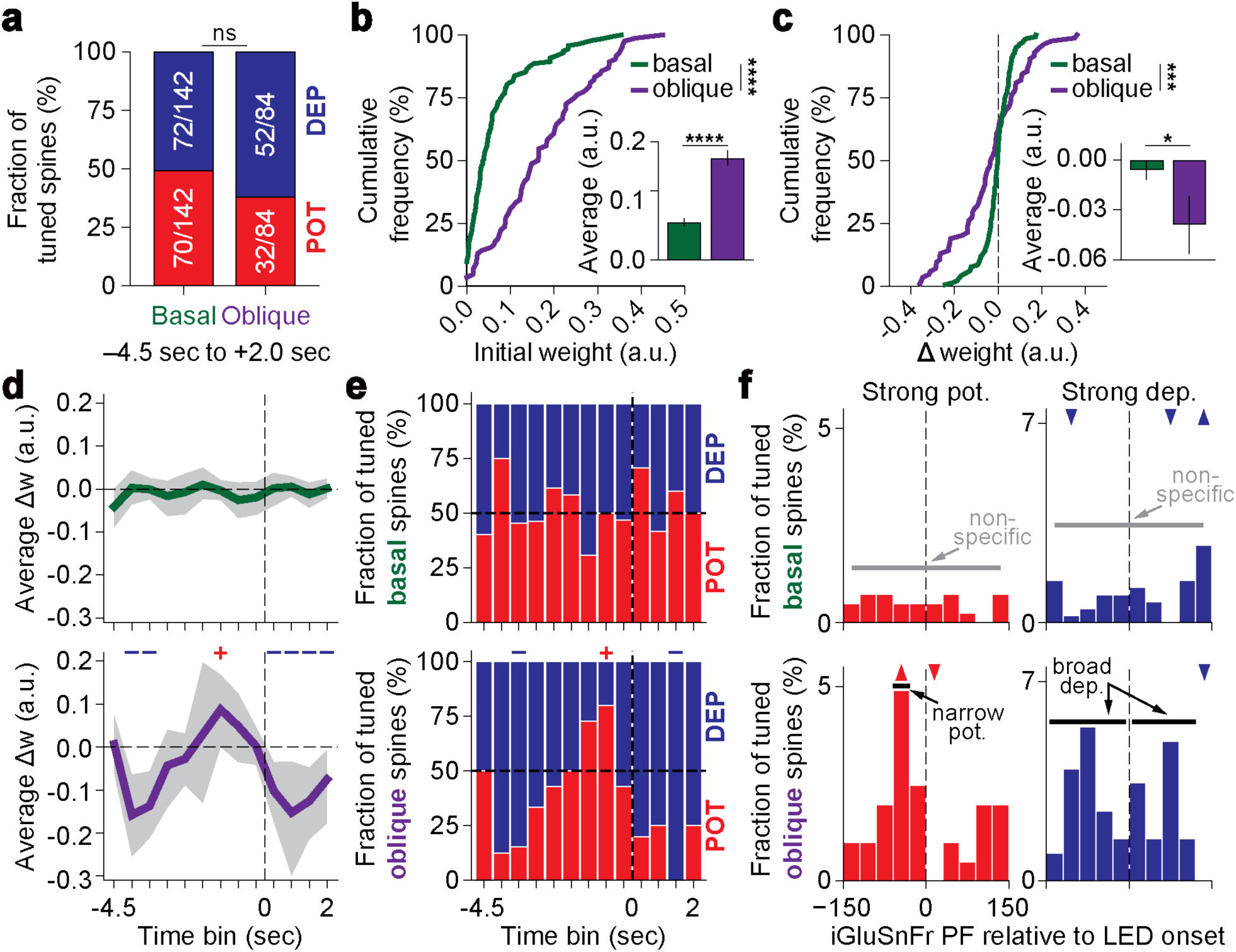
Compartment-specific expression of synaptic plasticity. **a**, Fraction of tuned basal and oblique spines undergoing potentiation (*Δw*>0) and depression (*Δw*≤0) around the time of optogenetic place field (PF) induction (–4.5 sec to +2.0 sec around LED onset; see Fig. 3b, left). Basal vs. Oblique, *P* = 0.1281 (Fisher’s exact test). **b**, Cumulative distribution of initial weights measured at tuned basal (*n* = 142 spines, green) and tuned oblique (n = 84 spines, purple) spines (Kolmogorov-Smirnov tests). Insets: average initial weight (a.u.) of all tuned basal and tuned oblique spines (mean ± s.e.m.), P < 0.0001 (Mann-Whitney two-tailed unpaired *t*-test). **c**, Cumulative distribution of plasticity events observed at tuned basal (*n* = 142 spines, green) and tuned oblique (n = 84 spines, purple) spines (Kolmogorov-Smirnov tests). Insets: average *Δw* (a.u.) of all tuned basal and tuned oblique spines (mean ± s.e.m.), P = 0.0476 (Mann-Whitney two-tailed unpaired *t*-test). **d**, Temporal profile of average plasticity occurring around the time of optogenetic PF induction (bin = 0.5 seconds; top: basal, *n* = 142 spines; bottom: oblique, *n* = 84 spines). Blue minus symbols depict time bins in which depression dominates, and red plus symbols depict those in which potentiation dominates (bootstrapped 95% confidence intervals (CIs) were generated from 10,000 bootstraps of the data points in each time bin; significance was assigned to time bins where CIs did not cover zero). **e**, Temporal profile of fraction of spines undergoing potentiation and depression around the time of optogenetic PF induction (bin = 0.5 seconds; top: basal, *n* = 142 spines; bottom: oblique, *n* = 84 spines). The blue minus (significant depression) and red plus (significant potentiation) symbols indicate time bins that lie outside the 95% CI generated from 1,000 data shuffles. **f**, Spatial profile of synaptic plasticity events (top row = basal spines; bottom row = oblique spines; bin = 30cm). Left: oblique spines receiving excitatory inputs tuned to locations before the stimulation site are selectively recruited to undergo strong potentiation, while oblique spines receiving excitatory inputs tuned to locations after the stimulation site are actively suppressed from undergoing potentiation. Right: oblique spines undergo broad depression around the induction site. Data were thresholded only to include the upper 25^th^ percentile of all plasticity events occurring at tuned spines. The up and down arrowheads indicate position bins that lie above or below the 95% CI generated from 1,000 data shuffles, respectively.

We further examined these findings by comparing the spatial and temporal kernels of changes in synaptic weights in each dendritic compartment. Notably, we observed that these profiles were exclusively expressed in spines located along oblique dendrites and completely absent in spines located along basal dendrites (Fig. 4d-f). These differences in dendritic domain-specific plasticity could not be attributed to artifactual differences in optogenetic stimulation or illumination of basal versus oblique dendrites that could have resulted from a gradual reduction in LED power along the basal-apical axis (Extended Data Fig. 4a, b). Our findings collectively demonstrate that the expression of synaptic plasticity is compartment-specific and is primarily driven by bidirectional changes in synaptic weights of inputs received by apical oblique rather than basal dendrites of CA1PNs *in vivo*.

## Discussion

In this study, we use an all-optical approach to monitor synaptic weight changes associated with *in vivo* PF formation in CA1PNs. We induce plasticity in hippocampal neurons on-demand in a manner largely identical to endogenous plasticity processes, using optogenetic techniques to trigger *de novo* PF formation in individual cells. This controlled and scalable approach allows for the targeted induction of plasticity in individual neurons and the dissection of links between synaptic plasticity and the emergence of spatial tuning in CA1PNs *in vivo*.

Our results provide important insights into the spatiotemporal organization of synaptic plasticity underlying feature selectivity in CA1PNs. First, we directly demonstrate a seconds-long, temporally-asymmetric synaptic plasticity kernel around induction stimulus, which was predicted by previous experimental and modeling studies of BTSP^16,17,23^: synaptic inputs active 1-2 seconds prior to the optogenetically induced depolarization are potentiated while synaptic inputs active outside this temporal window are depressed. We hypothesize that the primary mechanism of synaptic plasticity recorded in this study is the activity-dependent change in the postsynaptic abundance of AMPARs at synapses ^40–45^. Second, we demonstrate that properties of emergent PFs correlate with the number of synapses undergoing plasticity^16^. Third, we find that the site-specific potentiation of synaptic inputs occurring just prior to the location of PF induction are enveloped by a broad depression of excitatory synaptic inputs active outside this spatiotemporal window^23^. Finally, our results uncover an unexpected subcellular, dendritic compartment-specific, difference in the spatiotemporal expression of synaptic plasticity associated with BTSP between basal and oblique dendrites. We speculate that these compartment-specific differences in the expression of synaptic plasticity might be due to the structure of the presynaptic inputs arriving onto these dendritic compartments^36–38,46^, or differences in postsynaptic molecular signaling mechanisms, such as the properties of intracellular calcium release from internal stores, which we recently demonstrated to be significantly different between these two dendritic compartments in CA1PNs *in vivo* ^21^. In the context of BTSP, the precise molecular mechanisms linking plateau induction to the temporal kernel of synaptic plasticity revealed in this study remain to be determined^21,47–49^. Our approach provides a controlled experimental framework to dissect the molecular signaling mechanisms of synaptic plasticity underlying learning and memory in behaving animals.

## Methods

### Mice

All experiments were conducted in accordance with National Institutes of Health (NIH) guidelines and with the approval of the Columbia University Institutional Animal Care and Use Committee. All experiments were performed on 3–4 month-old C57Bl/6J non-transgenic male mice. Mice were kept in the vivarium on a reversed 12-hour light-dark cycle and were housed with 3–5 mice per cage (temperature, 22–23°C; humidity, 40%).

### Cannulas

Cannulas (dimensions = height, 1.5–1.9 mm; top length, 6.6 mm; base length, 3.1 mm; width, 3.0 mm) were 3D-printed using 316L stainless steel (InterPRO Additive Manufacturing Group, US). Glass windows of 3.0 mm diameter and 0.13–0.16 mm thickness containing a rectangular opening of 0.2 mm by 0.35 mm offset 0.3 mm from the window center were laser cut (Potomac Photonics, US) and attached to the metal cannulas using a UV-curable adhesive (Norland optical adhesive 81, Thorlabs, US). A 0.02 mm thick silicon membrane (Silex LTD, UK) was glued to the bottom of the glass window using a UV-curable silicone adhesive (5091 Nuva-SIL, Loctite, US) to protect the brain from exposure to the exterior environment. Cannulas were sterilized with 70% ethanol and washed with cortex buffer before surgery.

### Surgery

We placed male C57BL/6J mice, aged 8–10 weeks, under anesthesia and secured them in a heated stereotactic mount for surgery. We lubricated their eyes (Puralube ophthalmic ointment, Dechra, US), disinfected their scalps with alternating wipes of 140-proof ethanol and povidone-iodine solution, and administered subcutaneous injections of Meloxicam and Bupivacaine. After scalp removal, we cleaned the exposed skull with 140-proof ethanol, dried it with an air duster jet, and reattached the surrounding skin to the skull using a tissue adhesive (Vetbond, 3M, US). We also treated the skull surface with UV-curable adhesive (Optibond, Kerr, US) to enhance dental cement adhesion. We then created a craniotomy matching the cannula’s shape, centered at -2.3 mm AP, 1.6 mm ML relative to bregma. Upon bone and dura removal, we exposed the mediolateral (ML) axonal fibers overlying the hippocampus by aspirating the cortex and flushing it with ice-cold cortex buffer. We visually divided these fibers into thicker and thinner types, with the thinner ones lying directly above the anterior-posterior (AP) fibers. After removing all the ML fibers without disturbing the AP fibers, we inserted the cannula flush with the exposed tissue, pushing it down by 1.6 mm. We dried the surgical site, glued the cannula to the bone using a tissue adhesive (Vetbond, 3M, US), and firmly secured it with dental cement (Lang Dental, Contemporary Ortho-Jet™ (black liquid/powder), US), adding a titanium head post to the skull for head fixation. We monitored and provided analgesia (Meloxicam) postoperatively for three days.

### Plasmids

pCAG-jGCaMP7b was generated by subcloning a jGCaMP7b insert (from pGP-AAV-syn-jGCaMP7b-WPRE, Addgene #104489) into a pCAG backbone. pCAG-NES-jRGECO1a was generated by subcloning a NES-jRGECO1a insert (from pAAV-Syn-NES-jRGECO1a-WPRE, Addgene #100854) into a pCAG backbone. pCAG-SF-Venus-iGluSnFR-A184S was generated by subcloning a SF-Venus-iGluSnFR-A184S insert (from pAAV.CAG.SF-Venus-iGluSnFR.A184S, Addgene #106201) into a pCAG backbone. pAAV-CaMKIIa-bReaChes-mRuby3 was subcloned from pAAV-CaMKIIa-bReaChes-EYFP (gift from Karl Deisseroth) by excising the EYFP sequence and replacing it with a mRuby3 fragment (cloned from pAAV-CAG-mRuby3-WPRE, Addgene #107744). pCAG-bReaChes was generated by subcloning a bReaChes insert (from pAAV-CaMKIIa-bReaChes-EYFP, a gift from Karl Deisseroth) into a pCAG backbone. Subcloning was carried out using the In-Fusion cloning kit (Takara Bio). Endotoxin-free plasmids were purified using silica columns (NucleoBond Xtra Midi EF, Macherey-Nagel/Takara Bio, US).

### Single-cell electroporation

Long-taper borosilicate glass pipettes (G200-3, Warner Instruments, MA, 2.0 mm OD, 1.16 mm ID) were pulled (DMZ puller, Zeitz Instrumente Vertriebs GmbH, Germany) and filled with an electroporation solution, which consisted of the following: saline (diluted to 1X: 155 mM K-gluconate, 10 mM KCl, 10 mM HEPES, 4 mM KOH, pH 7.3, 316 mOsm), Alexa Fluor 488-hydrazide (diluted to 0.166-0.333 mM, Invitrogen), and plasmids (diluted to 50 ng/µL with endotoxin-free water). Electroporation solutions were filtered with 0.45 µm and 0.20 µm filters (Nalgene Syringe Filters, Cellulose Acetate, 4 mm, Thermo Scientific) before loading into the glass pipettes, which were then mounted on a micromanipulator (PatchStar, Scientifica, UK) angled at 32 degrees. A custom-made electronic circuit was used to switch between a patch-clamp amplifier head-stage (BVC-700A, Dagan Corporation, MN, US), which was used to monitor pipette resistance, and a digitally gated stimulus isolator (ISO-Flex, AMPI, Israel), which was used to deliver electroporating voltage pulses (−4 to -5 V, 100 Hz, 0.5 ms pulse width, 1 s duration). Both the patch-clamp amplifier and the stimulus isolator were controlled by a digitizer (Digidata 1550B, Molecular Devices, CA, US) using a stimulation protocol generated in Clampex (version 10.7; Molecular Devices, CA).

During the electroporation procedure, mice were placed under light anesthesia using isoflurane. Implants were then thoroughly washed with 1X PBS, and the slit was visually inspected to ensure no debris was stuck to the silicone membrane. Pipettes were then slowly lowered into the 1X PBS bath (tip resistance = 4-6 MΩ; positive pressure = 50-200 mBar) and subsequently guided to the top of the slit under 920 nm two-photon excitation with a 40X objective. Once at the slit, the pipettes were lowered in Z until contact was made with the silicone membrane, at which time the pipettes were rapidly advanced and pushed through the silicon membrane at 32 degrees. Successful penetration of the silicon membrane was assessed by an increase of 1-2 MΩ in tip resistance compared to the resistance measured in the 1X PBS bath. If tip resistance increased more than 2 MΩ, pipettes were retracted back into the bath and were unclogged by applying large positive pressure (>500 mBar) or large voltage pulses (−90 V, 100 Hz, 0.5 ms pulse width, 1 s duration). After successful membrane penetration, pipettes were gradually advanced diagonally toward the dorsal hippocampal CA1-area pyramidal layer, about 120–150 µm beneath the imaging window (tip resistance = 6-8 MΩ; positive pressure = 20-30 mBar). Once contact with a pyramidal cell was made and assessed visually by contact between the tip of the pipette and the soma and by a 1-1.5 MΩ increase in tip resistance, the pressure was dropped to 6-12 mBar and the cell was electroporated (−4 to -5 V, 100 Hz, 0.5 ms pulse width, 1 s duration). The pipette was kept on the soma for an additional 4-8 seconds to confirm that the Alexa 488 dye filled the soma and dendrites before carefully retracting to ensure the nucleus was not pulled out.

### Virtual-reality system

The VR system was designed as previously described^34,35^. Mice were head-fixed above a running wheel surrounded by 5 LCD computer monitors, each of which displayed a fraction of a VR scene that continuously updated as the position of the animal advanced. All VR scenes were designed using the Unity game engine. The following measures were taken to prevent LCD screen light from contaminating the PMTs: 1) the imaging objective and cannula implant were wrapped with back aluminum foil tape (T205-1.0, Thorlabs); 2) the LCD screens were set to their lowest brightness level; and 3) all VR scenes were devoid of red-containing pixels.

### Behavior training

After a 7-day recovery period from implant surgery, mice were habituated to handling and head fixation. After two days of acclimation and free running on the VR wheel, mice were water-restricted to 85-90% of their original body weight and were trained to navigate a 3 m virtual environment, then rewarded with a small volume of water (∼1-2 uL per reward; small rewards ensured that the animal did not satiate and disengage from the task before the end of the experiment) for running. After the mice reached the end of the virtual track, the screens were briefly blanked for a 1-2-second inter-trial interval (ITI), and the mice were immediately teleported back to the beginning of the track. The training period lasted approximately two weeks. Initially, rewards were randomly delivered to encourage running and gradually reduced to 1-2 fixed reward zones as the animal’s performance improved. For the induction experiments, the mice were exposed to a new 3 m virtual environment for two days before the experiment, and the locations of the fixed water rewards were randomly relocated. This exposure to a novel environment and the relocation of the two fixed rewards were implemented to increase the animal’s attention to the environment and enhance the likelihood of a successful induction.

### *In vivo* two-photon imaging and data preprocessing

All imaging was conducted using a custom-built two-photon 8 kHz resonant scanner using large aperture fluorescence collection optics (primary dichroic: 45.0 x 65.0 x 3.0 mm T865lpxrxt Chroma, US; secondary dichroic: 52.0 x 72.0 x 3.0 mm FF560-FDi02-t3 Semrock, US; green emission filter: 50.8 mm FF01-520/70 Semrock, US; red emission filter: 50.8 mm FF01-650/150 Semrock, US; GaAsP PMTs for green and red channels, PMT2101 Thorlabs, US) paired with a Nikon 16x water immersion, 0.8 NA, 3.0 mm working distance objective. Laser power was controlled with a Pockel cell (350-80LA modulator, 320RM 401 driver, Conoptics, US) and ranged from 20 to 100 mW (measured after the objective) depending on the imaging depth of the dendrite of interest. Image acquisition was controlled through commercial software (ScanImage, Vidriotech, US).

All imaging was performed in awake, fully trained running animals (> four weeks after surgery). Animals not exhibiting prolonged, consistent running were not used for experiments because motion artifacts induced by run-stop behavior made resolving out spines in the motion-corrected time averages challenging. SF-Venus-iGluSnFR-A184S/jRGECO1a/bReaChes expressing cells were imaged at 1020 nm (Chameleon Ultra II, Coherent, US). All frame scans lasted 1–2 minutes and were acquired at 120-240 Hz (40–60x optical zoom; 64– 128 lines/frame, 64–128 pixels/line) from soma and dendrites (basal and oblique). Dendritic health was monitored throughout the experiment. No signs of phototoxicity (i.e., dendrite swelling or blebbing) were observed.

Frame scans were motion-corrected, and fluorescence was extracted from segmented regions of interest (ROIs) using the SIMA software package^50^. Recordings were visually assessed for residual motion, and any data with uncorrected motion artifacts were discarded from further analysis. All ROIs were manually drawn on motion-corrected, maximum-intensity projected dendritic segments using the membrane-bound glutamate sensor, iGluSnFr, as the reference channel. For dendritic spines, ROIs were drawn along the boundary of the spine head, 1-2 pixels beyond the edge of detectable fluorescence. ROIs were also drawn on spines located on the z-axis of the dendritic shaft. Only spines tracked for the entire duration of the experiment were included in the analysis. Multiple large ROIs were drawn in the empty regions of the field of view to eliminate background. To correct for bleaching, fluorescent traces were fitted and normalized to a natural cubic spline curve. Movement-induced artifacts (defined here as a decrease in signal fluorescence to less than 50% of its bleaching normalized baseline and with a z-score less than -3) in fluorescence signals were blanked and excluded from the analysis. For the purposes of extrapolating spine input timing (detailed below), the resulting de-bleached, motion-stabilized SF-Venus-iGluSnFR-A184S signal was used for analysis. To measure spine synaptic weight (detailed below), the de-bleached, motion-stabilized jRGECO1a signals were first high-pass filtered (Butterworth, 2nd order zero-phase, 0.2 Hz cutoff). The resulting de-bleached, motion-stabilized high-pass filtered jRGECO1a signals were deconvolved to detect putative events by applying the OASIS software package^51^ using an AR1 model with a pre-computed signal decay constant of 300 ms (Ar_g = 0.97; event threshold = 0.3).

### Optogenetic place field induction

Optogenetic PF inductions were performed as previously described^19,21^. DNA plasmid constructs (Fig. 1g: pCAG-jGCaMP7b + pAAV-CaMKIIa-bReaChes-mRuby3; Fig. 2b: pCAG-SF-Venus-iGluSnFR-A184S + pCAG-NES-jRGECO1a + pCAG-bReaChes) were diluted to 50ng/µL and electroporated into single dorsal hippocampal CA1-area pyramidal neurons. Successfully electroporated cells were imaged 48-72h post electroporation. Cells were stimulated for 1 second at a randomly chosen location for five laps with an ultrafast high-power (∼30-40mW measured after the objective) collimated LED at 630 nm (Prizmatix, 630 nm). In the event of an evident preexisting somatic PF (assessed visually during baseline activity recordings), the LED stimulation zone was moved as far away as possible from the existing putative PF. To image somatic jGCaMP7b dynamics during the stimulation (Fig. 1g), the 630 nm LED light was passed through a dichroic mirror that allowed red light to pass into the objective and deflected green collected light to the green PMT. The red PMT was turned off for the duration of the stimulation experiment. Cells expressing jRGECO1a were not imaged during the stimulation period (Fig. 2b). Instead, somatic activity was recorded immediately before and after the induction experiment. LED stimulation light pulses were delivered using custom-built electronics, which were synchronized with the blanking signal of the imaging beam Pockels cell, allowing pulsed LED stimulation during the flyback periods of the Y-galvanometer. Following this wide-field optical stimulation protocol, the remainder of the imaging session was used to acquire postinduction somatic and dendritic data.

### Task structure

Before beginning all experiments, we pre-selected which dendrites to image. We randomly sampled dendrites per cell and chose segments that were parallel with the field of view and in which we could visibly see spines. We did not image the trunk or proximal somatic regions (< ∼50 μm from the soma) because spine density is low in these compartments. The number of dendrites we sampled per cell also depended on the animal’s behavior, to ensure that we could track all the selected dendrites for the duration of the experiment. If the animal was a good runner (i.e., > 130 laps in 1 hour), we imaged up to ∼ ten dendrites per cell. However, if it was a poor runner (i.e., < 50 laps in 1 hour), we limited our sampling to ∼2-4 dendrites per cell.

The experiment consisted of 3 phases: *Pre*, *Induction*, and *Post*.

- *Pre*: The goal of this phase was to sample the tuning preference of the soma (using jRGECO1a), map the spatiotemporal distribution of the synaptic inputs to spines (using iGluSnFr), and measure the spine’s initial weight (using jRGECO1a) (analysis detailed below). Because our recordings were high power and short in duration (∼1-2 minutes to avoid photodamage), we rerecorded our ROIs multiple times and concatenated the scans so that, in total, we had roughly ten laps worth of activity per ROI. We performed all analyses on the concatenated data.
- *Induction*: The goal of this phase was to induce a PF (detailed above). Only behavioral data was recorded.
- *Post*: The goal of this phase was to confirm that a somatic PF was induced (using jRGECO1a) and measure the final synaptic weights of all spines (using jRGECO1a). As in the *Pre* phase, where possible, given the animal was not satiated and disengaged from the task, we re-recorded the ROIs and concatenated the scans so that, in total, we had roughly ten laps worth of activity per ROI. We performed all analyses on the concatenated data.

### Spatial tuning curves and place field detection

Spatial tuning and PF detection analyses were performed as previously described^20,21,34,52^. In short, all spatial tuning analyses used either glutamate or calcium fluorescence traces to approximate activity over time. To generate spatial tuning curves, we divided the virtual track into 100 evenly spaced bins (3 cm) and, for each bin, calculated the average ΔF/F from frames in which the animal’s velocity was greater than or equal to 1.0 cm/s. After normalizing for animal occupancy (by averaging activity across all laps), we smoothed the resulting trace with a Gaussian kernel (*σ* = 9 cm) to obtain the spatial tuning curve.

To detect PFs, we generated a baseline distribution of spatial tuning curves for each neuron by randomly and independently shifting the activity on each lap a random distance in a circular manner. Following this, we recalculated the smoothed, lap-averaged tuning curve as detailed above. We repeated this process a thousand times, determining the 95^th^ percentile of baseline tuning values at each spatial bin, which served as the threshold for significant spatial tuning (*p* < 0.05). Areas of space where the true spatial tuning curve surpassed this baseline threshold were identified as potential PFs. We then measured the PF width by finding the distance between the points where the actual tuning curve first rose above and then dropped back below the threshold curve. In order to focus our analysis on compartments (i.e., somas or spines) with clearly defined firing fields, we set an additional criterion that there must be activity within the PF boundaries for a minimum of 20% of laps.

### Induced place field formation analysis

Optogenetically induced PFs were analyzed as previously described^21^. Briefly, the *Pre-* and *Post*-induction somatic calcium fluorescence traces (jRGECO1a ΔF/F) were used to generate spatial tuning curves as described above. Induction efficacy was calculated by measuring the maximum difference between the *Pre* and *Post* tuning curves inside the LED zone, a symmetric 15 cm radius window centered on LED onset. Cells were considered to be successfully induced if there was at least a 5% increase in somatic activity following optogenetic induction (Fig. 2b). Peri-formation velocity (Fig. 2b, Fig 3e) was defined as the mean velocity observed from 1.5 s before to 1.0 s after LED onset during the first induction lap.

### Extrapolating input timing analysis

This analysis was aimed at estimating the amount of time elapsed between the arrival of individual synaptic input (measured using iGluSnFr) and when a PF is induced (controlled experimentally by selecting the location of LED stimulation). To do this, we first mapped the spatial location of all inputs arriving at spines using the iGluSnFr activity profile collected during the *Pre* phase of the experiment. Using this information, we took the peak location of the detected spine iGluSnFr PF and the location of the LED onset, then measured the average amount of time it took the animal to travel between both locations. The average amount of time was calculated using the animal’s behavior during the five induction laps. We restricted this analysis only to spines receiving significantly spatially tuned synaptic input. Positive time values mean the input arrived after the LED onset. Negative time values mean the input arrived before the LED onset (Fig. 2c).

### Calculating synaptic weights analysis

All spines were assigned initial and final synaptic weights using the calcium (jRGECO1a) activity profile collected during the *Pre* and *Post* phases of the experiment. To measure synaptic weights, the amplitude of all isolated spine-specific calcium events were summed (OASIS-detection method described above). To isolate spine-specific activity, all spine events co-occurring with or active 0.1 seconds after a globally detected dendritic calcium event were removed from the analysis. Spines with no detected events or with no events remaining after the exclusion of global events were assigned an initial or final weight of 0. Changes in synaptic weight were calculated by measuring the difference (final weight – initial weight). Positive changes in synaptic weight mean the spine underwent potentiation, while negative changes in synaptic weight mean the spine underwent depression.

### Author Contribution

K.C.G., F.P., and A.L. conceived the study and designed the experiments; K.C.G. performed all the experiments; K.C.G. analyzed the data with the help of A.N. and Z.L; A.N. developed the electroporation protocol and built the resonant scanner microscope; K.C.G., F.P., and A.L. wrote the manuscript with input from all authors.

---

**Extended Data Fig. 1.**
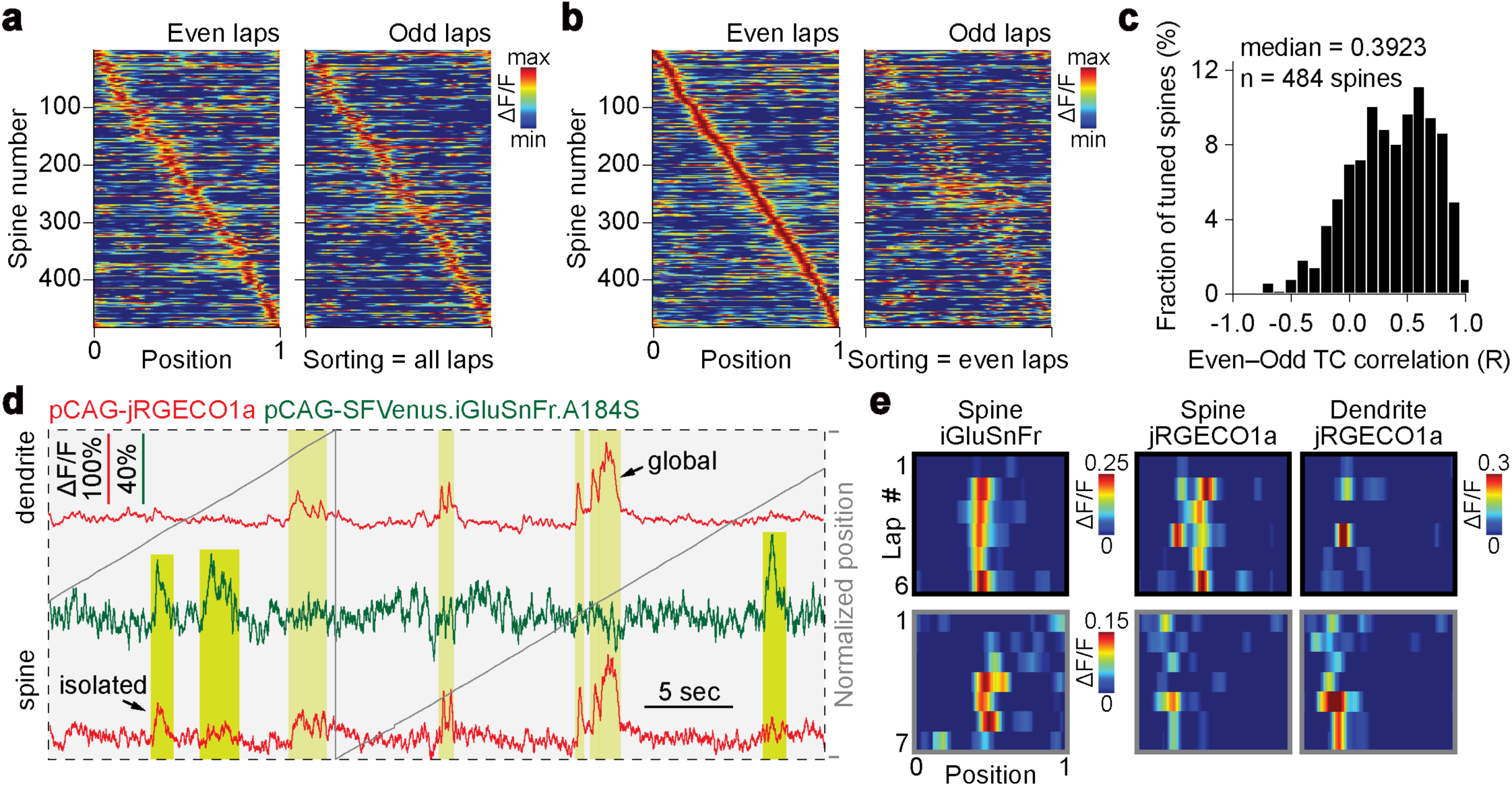
Spatial tuning of excitatory inputs. **a**, Average iGluSnFr ΔF/F across space for all tuned spines sorted by peak location (*n* = 484 spines from 15 induction experiments (12 cells)). Heat maps for even and odd laps are shown separately. Spines are ordered according to their peak location during all laps (see Fig. 2, panel f). **b**, Same as **a**, except spines are ordered according to their peak location during the even laps. **c**, Distribution of synaptic iGluSnFr even–odd lap correlations (median Pearson’s correlation coefficient = 0.3923). **d**, Example fluorescence traces of synaptic glutamate–calcium and dendritic calcium signals during navigation. Glutamate-evoked and non-evoked spine calcium responses are highlighted. Gray traces represent the animal’s position on the virtual track. **e**, Activity heatmaps for two spines receiving spatially tuned iGluSnFr input and associated spine and dendritic calcium activity profiles. Using spine calcium signals to extract input timing leads to erroneous time measurements because spine calcium signals can be strongly contaminated by global dendritic events (bottom row, gray). In other spines, using either spine glutamate or spine calcium signals leads to the correct measurement of input timing (top row, black).

**Extended Data Fig. 2.**
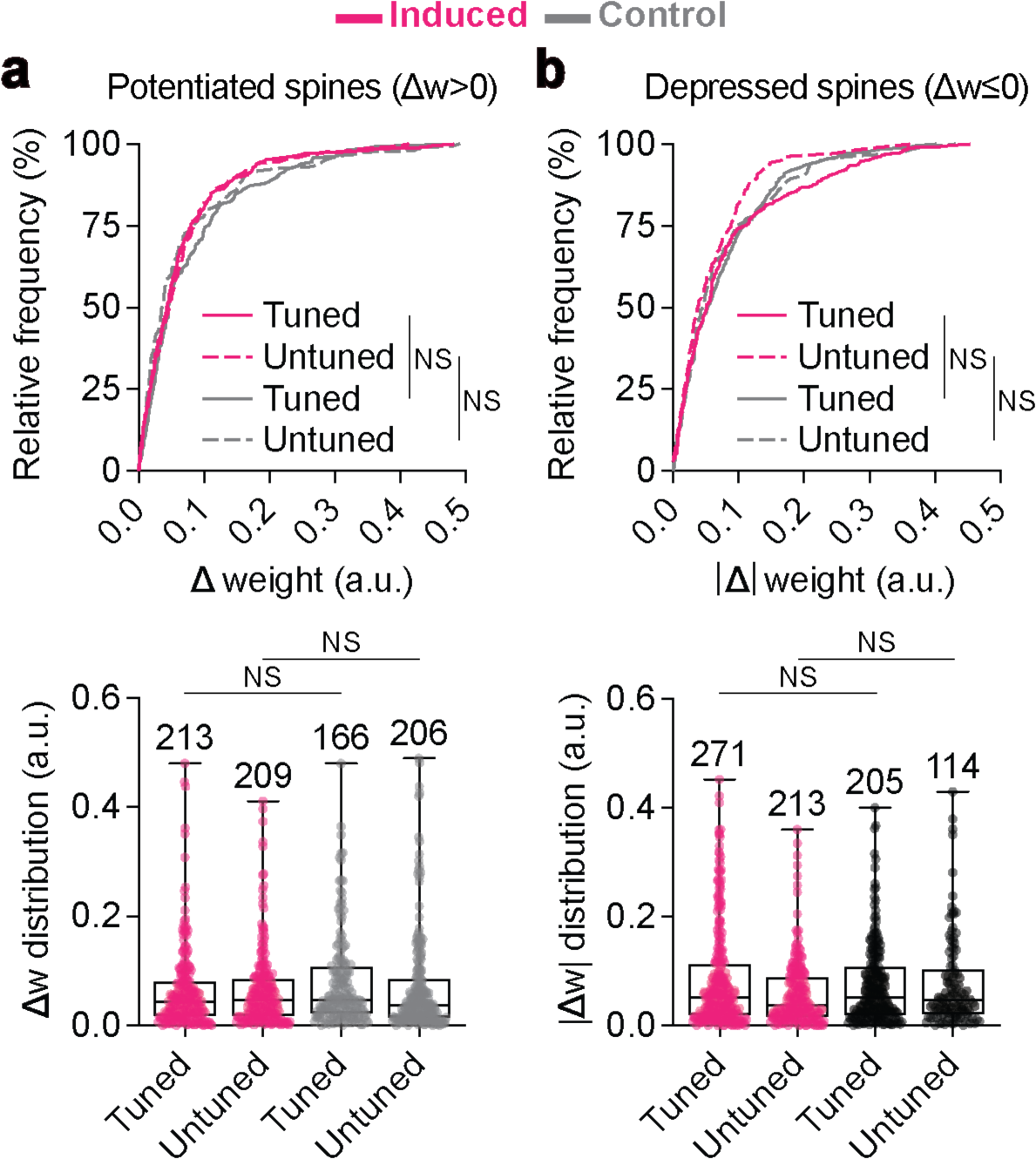
Similar magnitude and distribution of synaptic plasticity recorded in optogenetically induced place cells and non-induced cells. **a**, Top: Cumulative distribution of potentiation plasticity events observed at tuned and untuned spines of optogenetically induced place cells (magenta) and non-induced cells (gray) (Kolmogorov-Smirnov tests). Bottom: Quantification of the potentiation plasticity events occurring at tuned and untuned spines of optogenetically induced place cells (magenta) and non-induced cells (gray) (Kruskal-Wallis tests). **b**, Same as a, except looking at depression plasticity events.

**Extended Data Fig. 3.**
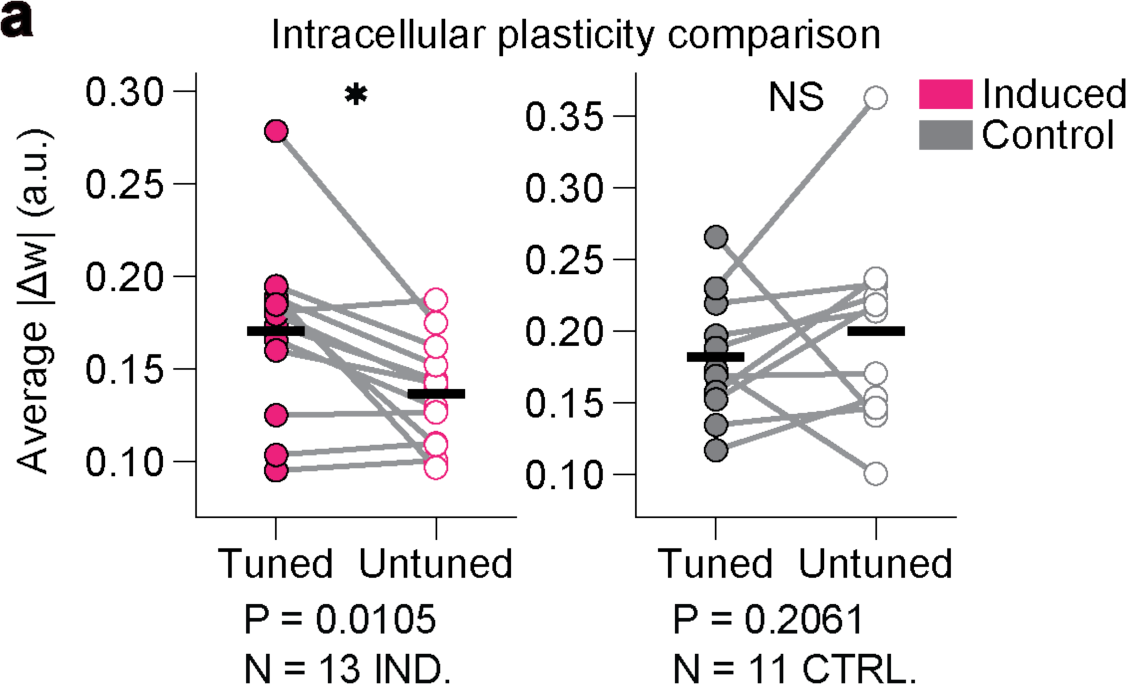
Place field induction selectively drives larger changes in synaptic weight at tuned spines compared to untuned spines. **a**, Average magnitude of plasticity event (regardless of direction, i.e., potentiation or depression) happening at tuned and untuned spines in induced (left, magenta) and non-induced (right, gray) cells (Wilcoxon matched-pairs signed rank two-tailed test: (Induced) *n* = 13 induction experiments, *P* = 0.0105; (Control) *n* = 11 control experiments, *P* = 0.2061)). Data were thresholded only to include the upper 25^th^ percentile of all plasticity events occurring at tuned and untuned spines.

**Extended Data Fig. 4.**
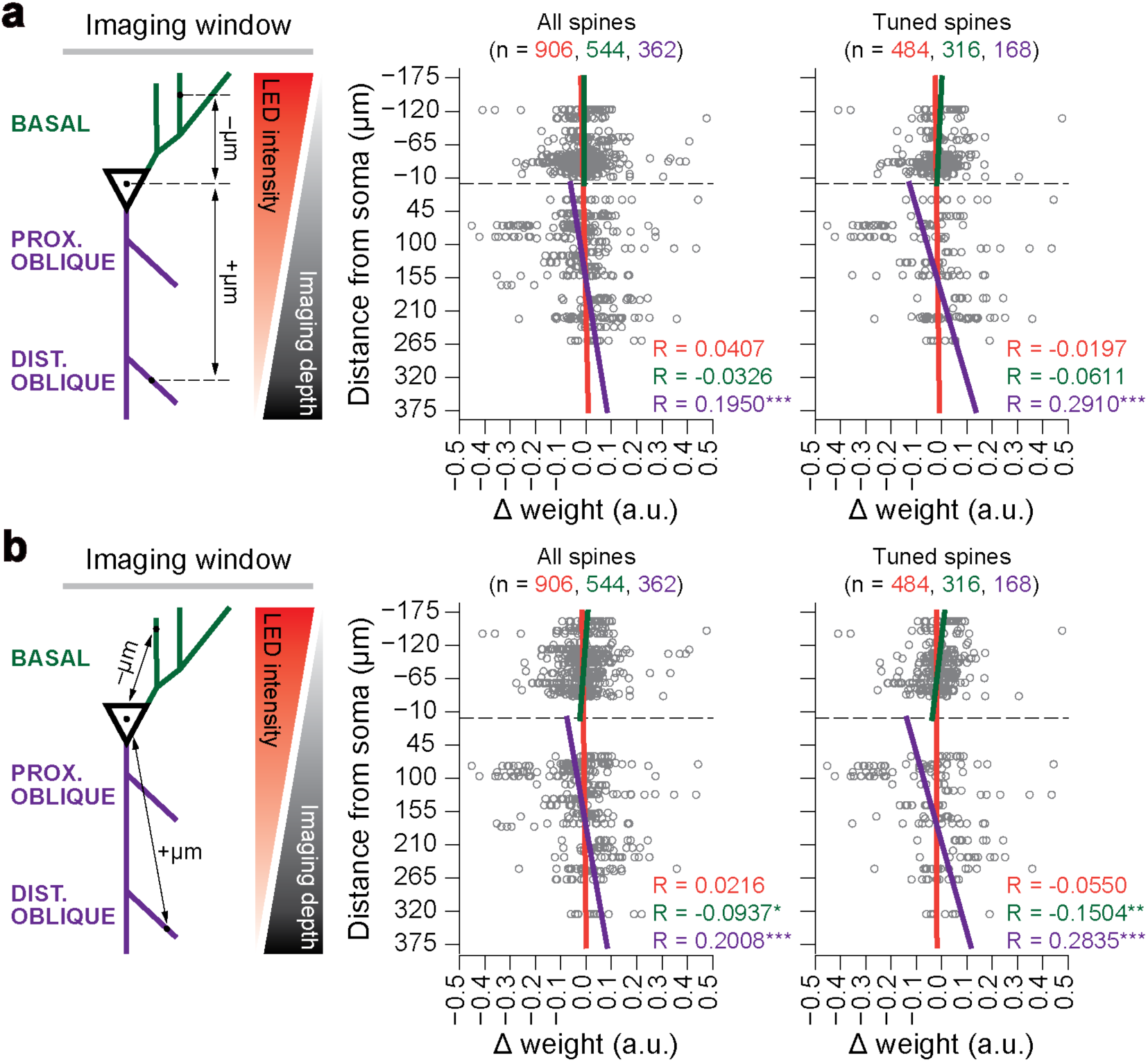
Differences in layer-specific plasticity cannot be explained by a gradient in LED power along the basal–apical axis. **a**, Left: Schematic illustrating method for measuring the distance between the soma and the recorded dendritic segment. We hypothesized that if the LED stimulations differentially excite basal and oblique dendrites due to a gradual reduction in power along the basal–apical axis, this would result in a plasticity gradient across all the layers. Center: Distribution of plasticity events observed at all spines of optogenetically induced place cells as a function of distance from the soma (Spearman r (all spines, orange): *n* = 906, two-tailed *t*-test, *R* = 0.0407, *P* = 0.2207; Spearman r (basal spines, green): *n* = 544, two-tailed *t*-test, *R* = –0.0326, *P* = 0.4476; Spearman r (oblique spines, purple): *n* = 362, two-tailed *t*-test, *R* = 0.1950, *P* = 0.0002). Right: Same as the center graph, except only looking at tuned spines (Spearman r (all spines, orange): *n* = 484, two-tailed *t*-test, *R* = –0.0197, *P* = 0.6652; Spearman r (basal spines, green): *n* = 316, two-tailed *t*-test, *R* = – 0.0611, *P* = 0.2787; Spearman r (oblique spines, purple): *n* = 168, two-tailed *t*-test, *R* = 0.2910, *P* = 0.0001). Data points fit by linear equation (all spines, orange; basal spines, green; oblique spines, purple). Spines located on the same dendritic branch were assigned the same distance from the soma. **b**, Same as a, except using a different method for measuring the distance between the soma and the recorded dendritic segment. Center: Distribution of plasticity events observed at all spines of optogenetically induced place cells as a function of distance from the soma (Spearman r (all spines, orange): *n* = 906, two-tailed *t*-test, *R* = 0.0216, *P* = 0.5158; Spearman r (basal spines, green): *n* = 544, two-tailed *t*-test, *R* = –0.0937, *P* = 0.0288; Spearman r (oblique spines, purple): *n* = 362, two-tailed *t*-test, *R* = 0.2008, *P* = 0.0001). Right: Same as the center graph, except only looking at tuned spines (Spearman r (all spines, orange): *n* = 484, two-tailed *t*-test, *R* = –0.0550, *P* = 0.2271; Spearman r (basal spines, green): *n* = 316, two-tailed *t*-test, *R* = –0.1504, *P* = 0.0074; Spearman r (oblique spines, purple): *n* = 168, two-tailed *t*-test, *R* = 0.2835, *P* = 0.0002). Data points fit by linear equation (all spines, orange; basal spines, green; oblique spines, purple). Spines located on the same dendritic branch were assigned the same distance from the soma.

**Extended Data Fig. 5.**
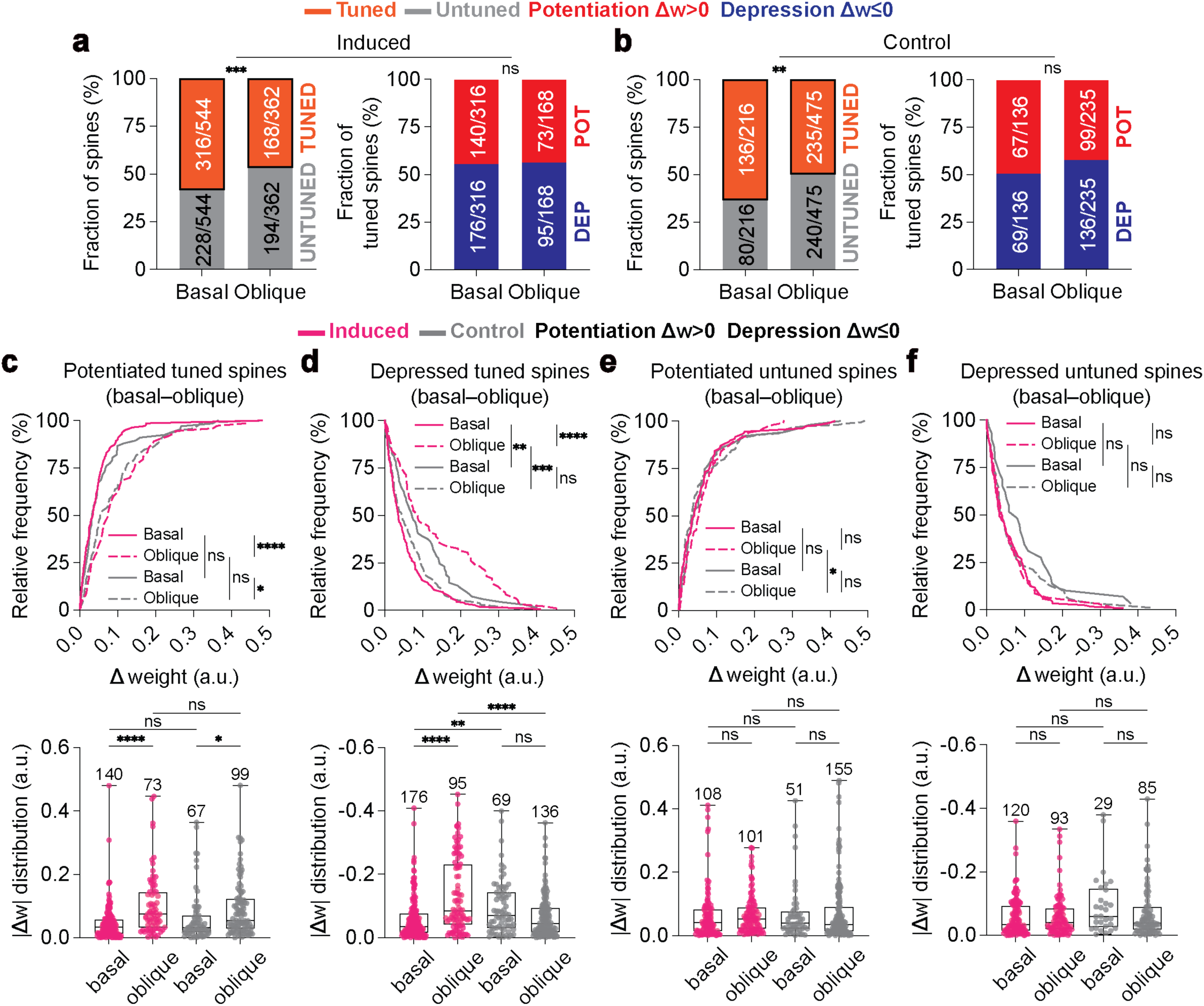
Intracellular comparison of spatial tuning properties and plasticity events occurring in basal and oblique spines of induced and non–induced cells. **a**, Induction experiments. Left: fraction of basal and oblique spines receiving spatially tuned synaptic iGluSnFr input (orange) and spines receiving no tuned input (gray). Basal vs. Oblique, *P* = 0.0007 (Fisher’s exact test). Right: fraction of tuned basal and oblique spines undergoing potentiation (*Δw*>0) and depression (*Δw*≤0). Basal vs. Oblique, *P* = 0.9234 (Fisher’s exact test). **b**, Control experiments. Left: fraction of basal and oblique spines receiving spatially tuned synaptic iGluSnFr input (orange) and spines receiving no tuned input (gray). Basal vs. Oblique, *P* = 0.0010 (Fisher’s exact test). Right: fraction of tuned basal and oblique spines undergoing potentiation (*Δw*>0) and depression (*Δw*≤0). Basal vs. Oblique, *P* = 0.1947 (Fisher’s exact test). **c**, Top: Cumulative distribution of potentiation plasticity events observed at tuned basal and oblique spines of optogenetically induced place cells (magenta) and non-induced cells (gray) (Kolmogorov-Smirnov tests). Bottom: Quantification of the potentiation plasticity events (Kruskal-Wallis tests). **d–f**, Same as **c**, except looking at depression plasticity events at tuned spines (**d**), potentiation plasticity events at untuned spines (**e**), and depression plasticity events at untuned spines (**f**). All box plots depict the median (central line) and interquartile range (25^th^ and 75^th^ percentile). Whiskers extend to the min-max data points.

**Extended Data Fig. 6.**
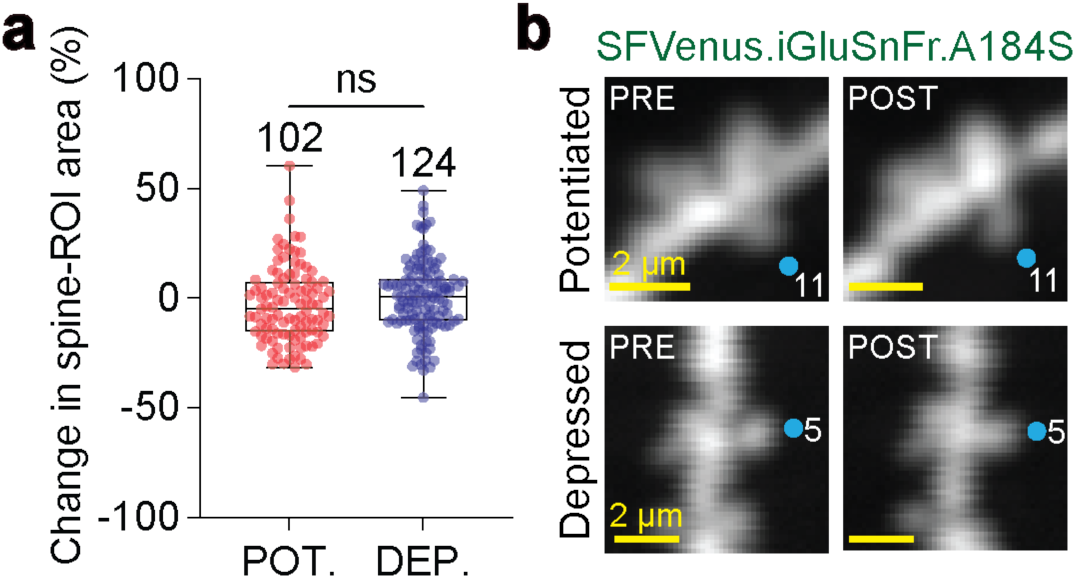
No detectable changes in spine head size between potentiated and depressed spines following place cell induction. **a**, Change in spine-ROI area of potentiated and depressed tuned spines active inside the plasticity kernel (−4.5 to +2.0 seconds). Percent change was calculated by measuring the difference in the area of the manually drawn ROI following induction (area after induction minus area before induction) divided by the area before induction (Mann-Whitney two-tailed unpaired *t*-test: Pot., *Δw*>0, n = 102 spines; Dep., *Δw*≤0, n = 124 spines; P = 0.0803). All box plots depict the median (central line) and interquartile range (25th and 75th percentile). Whiskers extend to the min-max data points. **b**, Magnified and cropped images of spines undergoing potentiation and depression (see Fig. 2d) but no detectable changes in spine head size. All ROIs were drawn on motion-corrected, maximum-intensity projected dendritic segments using the membrane-bound glutamate sensor, SFVenus.iGluSnFr.A184S, as the reference channel (see methods).

## References

1. Takeuchi, T., Duszkiewicz, A. J. & Morris, R. G. M. The synaptic plasticity and memory hypothesis: encoding, storage and persistence. Philosophical Transactions of the Royal Society B: Biological Sciences vol. 369 20130288 Preprint at 10.1098/rstb.2013.0288 (2014).

2. Magee, J. C. & Grienberger, C. Synaptic Plasticity Forms and Functions. Annu. Rev. Neurosci. 43, 95–117 (2020).

3. Bliss, T. V. & Lomo, T. Long-lasting potentiation of synaptic transmission in the dentate area of the anaesthetized rabbit following stimulation of the perforant path. J. Physiol. 232, 331–356 (1973).

4. Bliss, T. V. & Collingridge, G. L. A synaptic model of memory: long-term potentiation in the hippocampus. Nature 361, 31–39 (1993).

5. Malenka, R. C. & Bear, M. F. LTP and LTD: an embarrassment of riches. Neuron 44, 5–21 (2004).

6. O’Keefe, J. & Dostrovsky, J. The hippocampus as a spatial map. Preliminary evidence from unit activity in the freely-moving rat. Brain Res. 34, 171–175 (1971).

7. Tolman, E. C. Cognitive maps in rats and men. Psychol. Rev. 55, 189–208 (1948).

8. Moser, E. I., Kropff, E. & Moser, M.-B. Place cells, grid cells, and the brain’s spatial representation system. Annu. Rev. Neurosci. 31, 69–89 (2008).

9. Keefe, J. & Nadel, L. The Hippocampus As a Cognitive Map. (Clarendon Press, 1978).

10. Scoville, W. B. & Milner, B. Loss of recent memory after bilateral hippocampal lesions. J. Neurol. Neurosurg. Psychiatry 20, 11–21 (1957).

11. Eichenbaum, H. A cortical-hippocampal system for declarative memory. Nat. Rev. Neurosci. 1, 41–50 (2000).

12. Squire, L. R. & Wixted, J. T. The cognitive neuroscience of human memory since H.M. Annu. Rev. Neurosci. 34, 259–288 (2011).

13. Epsztein, J., Brecht, M. & Lee, A. K. Intracellular determinants of hippocampal CA1 place and silent cell activity in a novel environment. Neuron 70, 109–120 (2011).

14. Harvey, C. D., Collman, F., Dombeck, D. A. & Tank, D. W. Intracellular dynamics of hippocampal place cells during virtual navigation. Nature 461, 941–946 (2009).

15. Lee, D., Lin, B.-J. & Lee, A. K. Hippocampal place fields emerge upon single-cell manipulation of excitability during behavior. Science 337, 849–853 (2012).

16. Bittner, K. C., Milstein, A. D., Grienberger, C., Romani, S. & Magee, J. C. Behavioral time scale synaptic plasticity underlies CA1 place fields. Science 357, 1033–1036 (2017).

17. Bittner, K. C. et al. Conjunctive input processing drives feature selectivity in hippocampal CA1 neurons. Nat. Neurosci. 18, 1133–1142 (2015).

18. Fan, L. Z. et al. All-optical physiology resolves a synaptic basis for behavioral timescale plasticity. Cell 186, 543–559.e19 (2023).

19. Rolotti, S. V. et al. Local feedback inhibition tightly controls rapid formation of hippocampal place fields. Neuron 110, 783–794.e6 (2022).

20. Geiller, T. et al. Local circuit amplification of spatial selectivity in the hippocampus. Nature 601, 105–109 (2022).

21. O’Hare, J. K. et al. Compartment-specific tuning of dendritic feature selectivity by intracellular Ca2+ release. Science 375, eabm1670 (2022).

22. Zhao, X., Wang, Y., Spruston, N. & Magee, J. C. Membrane potential dynamics underlying context-dependent sensory responses in the hippocampus. Nat. Neurosci. 23, 881–891 (2020).

23. Milstein, A. D. et al. Bidirectional synaptic plasticity rapidly modifies hippocampal representations. Elife 10, (2021).

24. Marvin, J. S. et al. Stability, affinity, and chromatic variants of the glutamate sensor iGluSnFR. Nat. Methods 15, 936–939 (2018).

25. Adoff, M. D. et al. The functional organization of excitatory synaptic input to place cells. Nat. Commun. 12, 3558 (2021).

26. Chen, W. et al. In vivo volumetric imaging of calcium and glutamate activity at synapses with high spatiotemporal resolution. Nat. Commun. 12, 6630 (2021).

27. Higley, M. J. & Sabatini, B. L. Calcium signaling in dendritic spines. Cold Spring Harb. Perspect. Biol. 4, a005686–a005686 (2012).

28. Bloodgood, B. L. & Sabatini, B. L. Ca2+ signaling in dendritic spines. Curr. Opin. Neurobiol. 17, 345–351 (2007).

29. Bloodgood, B. L., Giessel, A. J. & Sabatini, B. L. Biphasic synaptic ca influx arising from compartmentalized electrical signals in dendritic spines. PLoS Biol. 7, e1000190 (2009).

30. Harnett, M. T., Makara, J. K., Spruston, N., Kath, W. L. & Magee, J. C. Synaptic amplification by dendritic spines enhances input cooperativity. Nature 491, 599–602 (2012).

31. Weber, J. P. et al. Location-dependent synaptic plasticity rules by dendritic spine cooperativity. Nat. Commun. 7, (2016).

32. Higley, M. J. & Sabatini, B. L. Calcium signaling in dendrites and spines: Practical and functional considerations. Neuron 59, 902–913 (2008).

33. Siegel, F. & Lohmann, C. Probing synaptic function in dendrites with calcium imaging. Exp. Neurol. 242, 27–32 (2013).

34. Priestley, J. B., Bowler, J. C., Rolotti, S. V., Fusi, S. & Losonczy, A. Signatures of rapid plasticity in hippocampal CA1 representations during novel experiences. Neuron 110, 1978–1992.e6 (2022).

35. Bowler, J. C. & Losonczy, A. Direct cortical inputs to hippocampal area CA1 transmit complementary signals for goal-directed navigation. bioRxiv (2022) doi:10.1101/2022.11.10.516009.

36. Rigby, M. et al. Multi-synaptic boutons are a feature of CA1 hippocampal connections in the stratum oriens. Cell Rep. 42, 112397 (2023).

37. Li, X. G., Somogyi, P., Ylinen, A. & Buzsáki, G. The hippocampal CA3 network: an in vivo intracellular labeling study. J. Comp. Neurol. 339, 181–208 (1994).

38. Grillo, F. W. et al. A distance-dependent distribution of presynaptic boutons tunes frequency-dependent dendritic integration. Neuron 99, 275–282.e3 (2018).

39. Soltesz, I. & Losonczy, A. CA1 pyramidal cell diversity enabling parallel information processing in the hippocampus. Nat. Neurosci. 21, 484–493 (2018).

40. Shi, S.-H. et al. Rapid spine delivery and redistribution of AMPA receptors after synaptic NMDA receptor activation. Science 284, 1811–1816 (1999).

41. Hayashi, Y. et al. Driving AMPA receptors into synapses by LTP and CaMKII: Requirement for GluR1 and PDZ domain interaction. Science 287, 2262–2267 (2000).

42. Zhang, Y., Cudmore, R. H., Lin, D.-T., Linden, D. J. & Huganir, R. L. Visualization of NMDA receptor-dependent AMPA receptor synaptic plasticity in vivo. Nat. Neurosci. 18, 402–407 (2015).

43. Groc, L. & Choquet, D. Linking glutamate receptor movements and synapse function. Science 368, (2020).

44. Borgdorff, A. J. & Choquet, D. Regulation of AMPA receptor lateral movements. Nature 417, 649–653 (2002).

45. Huganir, R. L. & Nicoll, R. A. AMPARs and synaptic plasticity: the last 25 years. Neuron 80, 704–717 (2013).

46. Takács, V. T., Klausberger, T., Somogyi, P., Freund, T. F. & Gulyás, A. I. Extrinsic and local glutamatergic inputs of the rat hippocampal CA1 area differentially innervate pyramidal cells and interneurons. Hippocampus 22, 1379–1391 (2012).

47. Xiao, K., Li, Y., Chitwood, R. A. & Magee, J. C. A critical role for CaMKII in behavioral timescale synaptic plasticity in hippocampal CA1 pyramidal neurons. Sci. Adv. 9, (2023).

48. Caya-Bissonnette, L., Naud, R. & Béïque, J.-C. Cellular substrate of eligibility traces. bioRxiv (2023) doi:10.1101/2023.06.29.547097.

49. Moldwin, T., Azran, L. S. & Segev, I. A generalized framework for the calcium control hypothesis describes weight-dependent synaptic changes in behavioral time scale plasticity. bioRxiv (2023) doi:10.1101/2023.07.13.548837.

50. Kaifosh, P., Zaremba, J. D., Danielson, N. B. & Losonczy, A. SIMA: Python software for analysis of dynamic fluorescence imaging data. Front. Neuroinform. 8, 80 (2014).

51. Friedrich, J., Zhou, P. & Paninski, L. Fast online deconvolution of calcium imaging data. PLoS Comput. Biol. 13, e1005423 (2017).

52. Geiller, T. et al. Large-scale 3D two-photon imaging of molecularly identified CA1 interneuron dynamics in behaving mice. Neuron 108, 968–983.e9 (2020).

